# A Screen To Identify Protein Phosphatases with Roles in Circadian Period, Temperature Compensation and Output in the *Neurospora* Circadian Clock

**DOI:** 10.64898/2026.05.20.723021

**Authors:** Adrienne K. Mehalow, Bin Wang, Jennifer J. Loros, Jay C. Dunlap

## Abstract

The circadian clock is a highly conserved evolutionary advantage which allows organisms to anticipate regular changes in daily environmental conditions. Clocks from fungi to mammals rely on a transcription-translation feedback loop (TTFL) mechanism. Phosphorylation is understood to be a critical regulatory step for maintaining the period of the circadian clock and feedback loop closure. The role of kinases in the Neurospora clock has been examined extensively; however, phosphatases have not been systematically interrogated. By re-examining the Neurospora genome using current informatic tools we identified the 30 genes previously identified as encoding protein phosphatases as well as 13 novel genes, and we assessed the function of the core circadian clock in 39 non-essential phosphatases using a real-time luciferase reporter. We observed both period lengthening and shortening effects, which are not restricted to a single phosphatase family or fold. All but one deletion mutant maintained a rhythmic core clock. In addition, we observed a new temperature compensation defect in the previously studied knockout of phosphatase *pph-4*, the result of nutritional growth conditions.

## INTRODUCTION

Circadian biology is the study of oscillating biological systems which exhibit a period of approximately 24 hours to align with the predictable day/night cycle of life on Earth[1]. Nearly all organisms on the planet contain an intrinsic circadian clock which allows the anticipation of daily environmental changes and confers a selective advantage. While the individual clock proteins are not conserved across the tree of life, the clock architecture of a core transcription- translation-feedback-loop (TTFL) driving diverse circadian outputs appears from fungi and plants all the way to mammals[2]. We use the fungal model organism *Neurospora crassa* to interrogate regulation of the TTFL and outputs from the circadian clock[3]. *Neurospora* is readily cultured in the haploid stage of its lifecycle, is amenable to further genetic manipulations, and has a relatively small genome of ∼10,000 genes[4]. The *Neurospora* core clock consists of the positive acting heterodimeric transcription factor (TF) White Collar Complex (WCC), composed of the white collar 1 (WC-1) and white collar 2 (WC-2) proteins, and the Frequency protein (FRQ) acts as a scaffold to assemble a repressive complex, the FFC, that also contains FRQ-Interacting RNA Helicase (FRH) and Casein Kinase 1a (CK-1a) [5]. In brief, during the night FFC repressor activity reaches a nadir due to protein inactivation and degradation. In the absence of functional repressor, WCC complex becomes increasingly active and drives new transcription of *frq* mRNA. Peak WCC activity occurs at or near dawn, after which TF function is reduced by the rising level of active FFC repressor. The daily cycle is complete when onset of darkness again reduces the abundance of active repressor and allows WCC to engage in new mRNA synthesis.

Three of the core *Neurospora* clock proteins WC-1, WC-2, and FRQ, are further regulated by extensive post-translational modifications (PTMs). The best studied PTM in the context of the circadian clock is phosphorylation[6-8]. All three core clock proteins are nascently devoid of phosphorylation and become progressively modified in a regular daily paaern. Scores of phosphorylations have been identified on each of the three proteins[6, 7, 9]. While the phospho-code is complex and not fully understood, we know that some phosphorylation states must confer protein activity, while others specify inactivation. Multiple kinases, including CK1, CK2, CHK2 and GSK, are known to alter behavior of the circadian clock[10-13] but the principal ones are CK1 and CK2, and exhaus3ve screens of Neurospora kinases have failed to identify other kinases whose loss confers significant clock effects [14]. Because phosphatases oppose the activity of kinases, phosphatase mutants should also yield circadian phenotypes and indeed, several phosphatases have been implicated in circadian clock regulation. The best studied of these in *Neurospora* are PP1, PP2A, and PP4[15, 16]. However, many phosphatases in *Neurospora* remain unannotated and thus unstudied.

In this screen we report a more complete tree of *Neurospora* phosphatases identified with the bioinformatics tool HMMER[17]. We select phosphatases expected to target protein substrates, then conduct a reverse genetic screen for defects in circadian period, phase, temperature compensation, and output for the 39 non-essential enzymes. Our screen identifies both known and novel circadian effects aaributed to loss of individual phosphatases, and complements phosphatase screens performed in other organisms[18].

## RESULTS

We applied a previously published HMMER profile to identify unannotated phosphatases in the *Neurospora crassa* genome[19]. Our search yielded 408 accession matches corresponding to 93 unique genes (Supp Table1). Where multiple accessions for a gene matched the HMMER profile, the accession generating the smallest single-domain E-value was selected. E-values smaller than 1×10*-4 were assigned as high confidence hits, E-values between 1×10*-1 and 1×10*-4 were assigned as low confidence, and all larger E-values were considered false hits. Our final identification contained 78 putative phosphatases, representing nine of the ten folds described for human phosphatases[19] (Fig. 1, Supp. Fig. 1). Similar to yeast, only the Protein Histidine Phosphatase (PHP) fold was absent. This new screen identified all 30 of the *Neurospora* phosphatase genes previously identified [20] as well as 13 novel genes.

**Figure 1.**
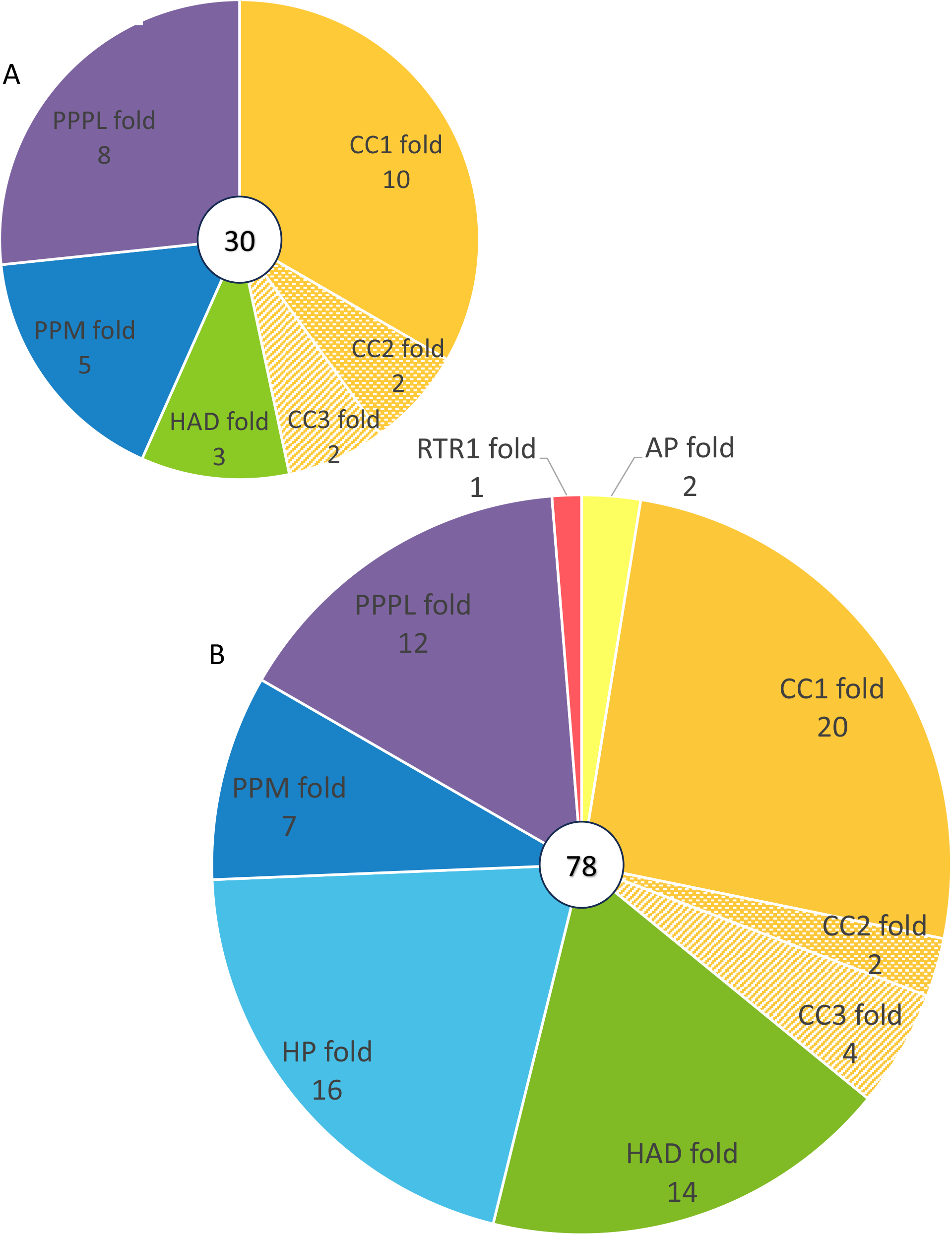
Comparison of phosphatases previously described in *Neurospora crassa* with those identified by our HMMER search. (A) Protein phosphatases published by Ghosh et al. assigned to their relevant phosphatase folds. (B) The 78 phosphatases identified in our HMMER search and their assigned folds. Protein and non-protein phosphatases are included. HP, RTR1, and AP folds previously had no annotated members in Neurospora.

While the total number of *Neurospora* phosphatases is very similar to yeast, the distribution of among folds and families reflects evolutionary changes. The CC1, PPPL, and HAD folds account of the majority of changes compared to the yeast phosphatome. The CC1 fold shows a decrease in the number of DSP (3 fewer members) family phosphatases, but an expansion in both the PTP (5 additional members) and PTEN (3 additional members) families compared to yeast. In contrast, the CC2 and CC3 folds are relatively unchanged. The OCA family, which is also present in Dictyostelium, yeast, and sponges, is entirely lost making *Neurospora* more similar to humans which also lack this family. The PPPL fold of *Neurospora* shows a loss of PPP family members compared to yeast (4 fewer members). This fold is highly conserved across species and believed to be the oldest branch of phosphatase evolution. Finally, the HAD fold of *Neurospora* is expanded, but only in the NagD family (5 additional members) of phosphatases. NagD family phosphatases have a broad range of substrates and are known to act also on nucleotide monophosphates and sugars. The remaining catalytic folds contained only minor changes in the total count of phosphatase members relative to yeast. Notably, our HMMER search assigns 16 *Neurospora* genes to the Histidine Phosphatase (HP) fold, which previously had no members identified in *Neurospora*.

Our interest is in identifying phosphatases which regulate period, temperature compensation (TC), phase, or output of the circadian clock. Because we know the circadian clock is controlled by Ser/Thr/Tyr phosphorylation events on WCC and FRQ proteins, we chose to focus on only on protein phosphatases. This allowed us to exclude some folds and sub- families from our circadian screen. For example, all 16 members of the (HP) fold were excluded because there is no evidence of histidine phosphorylation regulating any eukaryotic circadian clock. Similarly, the 5 members of the CC1-Sac family were excluded because all reported substrates are phosphatidyl-inositol (PI) species. We also excluded specific genes based on reported structure or function. For example, 2 arsenate reductases in the CC3-CDC25 family and the pseudo-phosphatase TIM50 in the HAD-FCP family were not screened. The final list selected for circadian screening contained 43 genes, composed of 30 originally published protein phosphatases and 1ti novel phosphatases from our HMMER search (Fig2).

**Figure 2.**
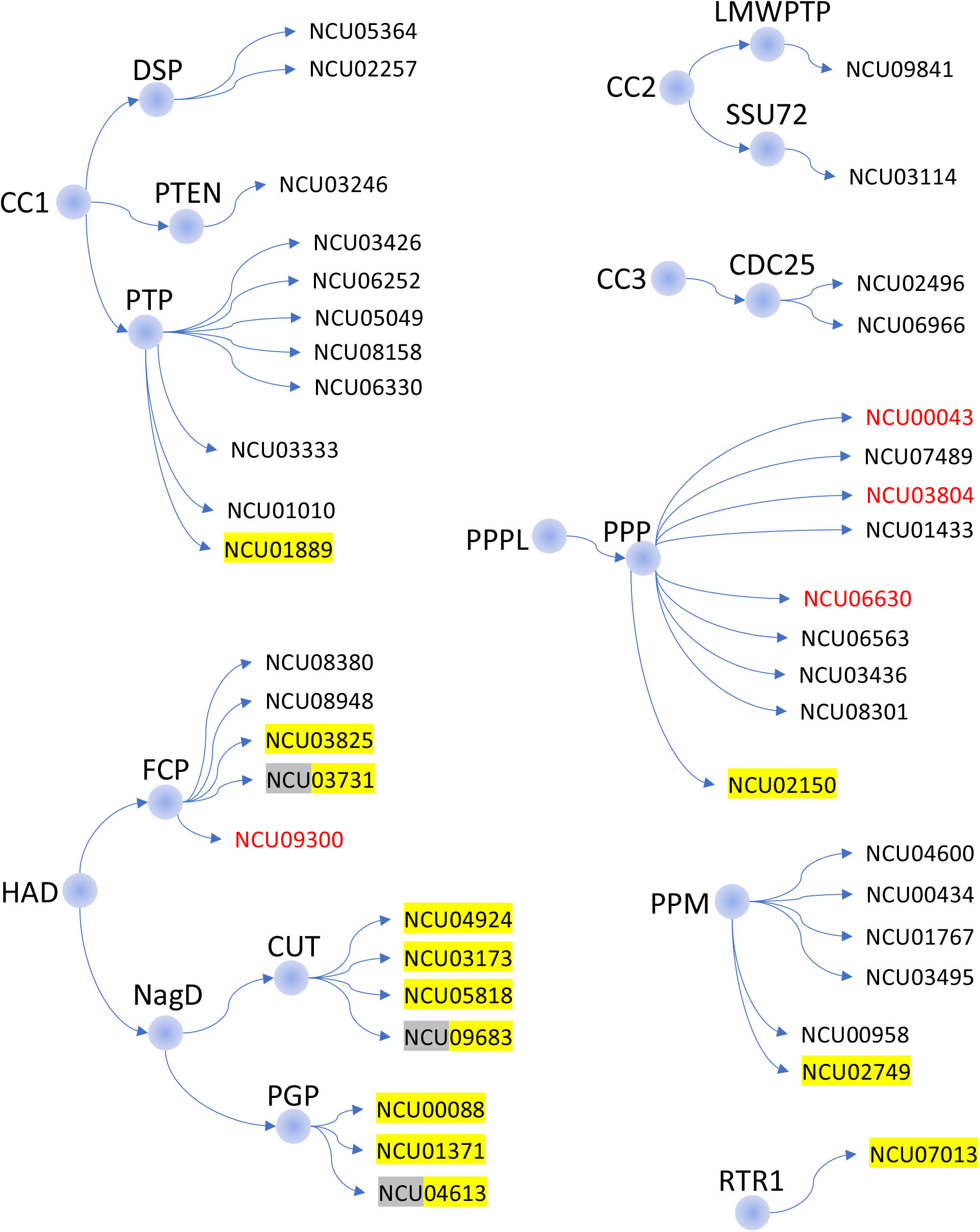
Protein phosphatases selected for the circadian screen, as assigned to folds and families by HMMER. More closely related phosphatases are placed next to each other; however, lines are not drawn to evolutionary scale. The four essential phosphatases (not screened) are in red. The 13 likely protein phosphatases not identified by Ghosh et al. are highlighted in yellow. Gray shading indicates genes which were low confidence matches to the HMM profile but were still included in the screen.

Of the 43 total phosphatases, 39 are non-essential which allowed us to easily screen gene knockouts for alterations in circadian period, phase, temperature compensation, and output (Table1). To our surprise, nearly all knockouts retained a functioning clock. We observe new long and short period mutants, phase delays, and a previously unappreciated temperature compensation defect among the knockouts.

**Table 1:** Circadian period, output, and temperature compensation for selected *Neurospora* phosphatases. Phosphatases are arranged in order of ascending period length, as measured by the core clock reporter. FGSC reference numbers are included for deletions derived from the *Neurospora* knockout collection. Fold and family are listed, as assigned by HMMER. Gene names and descriptions were obtained from FungiDB https://fungidb.org/app. Period of the core clock was determined over multiple runs and temperatures to produce temperature compensation curves. The calculated 25°C period and the Q10 value for each deletion are derived from the best-fit linear mixed model generated from all data for the strain.

| NCU | FGSC origin | Fold-Family-SubFamily (HMMER) | Symbol (FungiDb) | Description (FungiDb) | Core clock period @25C calc (hrs) | Core clock Q10 | Output period @25C (hrs) |
| --- | --- | --- | --- | --- | --- | --- | --- |
| N/A | WT2489 | N/A | N/A | Wildtype strain | 20.85 | 0.99 | 22.3 +/-0.4 |
| NCU08301 | 12453 and 12454 | PPPL-PPP | pph-4 | calcineurin-like phosphoesterase | 17.03 | 0.84 | arr |
| NCU04600 | 11185 | PPM | pph-9 | protein phosphatase-9 | 18.90 | 1.02 | 21.8 +/-0.4 |
| NCU01371 | 21200 | HAD-NagD-PGP | n/a | UPF0660 protein | 19.28 | 1.01 | arr |
| NCU00088 | 16344 | HAD-NagD-PGP | npp-1 | nitrophenylphosphatase-1 | 19.57 | 0.93 | 21.7 +/-0.3 |
| NCU02496 | 16180 | CC3-CDC25 | div-12 | M-phase inducer phosphatase 3 | 19.92 | 1.03 | nd |
| NCU09683 | 18536 | HAD-NagD-CUT | n/a | hypothetical protein | 20.19 | 0.97 | 21.6 +/-0.3 |
| NCU03173 | 18749 | HAD-NagD-CUT | n/a | HAD superfamily hydrolase | 20.32 | 1.00 | 22.2 +/-0.2 |
| NCU02749 | 19811 | PPM | n/a | hypothetical protein | 20.39 | 0.98 | 22.3 +/-0.2 |
| NCU04613 | 15152 | HAD-NagD-PGP | n/a | HAD superfamily hydrolase | 20.41 | 1.00 | 22.6 +/-0.2 |
| NCU04924 | 20352 | HAD-NagD-CUT | cut-1 | HAD-superfamily hydrolase | 20.42 | 0.98 | 21.2 +/-0.5 |
| NCU03731 | 18653 | HAD-FCP | n/a | HAD superfamily hydrolase | 20.48 | 0.98 | 22.1 +/-0.2 |
| NCU05818 | 20134 | HAD-NagD-CUT | n/a | Phosphatidyl synthase | 20.55 | 0.99 | 22.2 +/-0.1 |
| NCU02150 | 19090 | PPPL-PPP | stp-1 | serine-threonine protein phosphatase-1 | 20.55 | 1.00 | 21.9 +/-0.3 |
| NCU08948 | 18439 | HAD-FCP | pph-11 | NIF domain-containing protein | 20.57 | 0.95 | fc |
| NCU00958 | 19378 | PPM | azr-1 | azacytidine resistance-1 | 20.60 | 0.98 | 22.7 +/-0.2 |
| NCU09841 | 18801 | CC2-LMWPTP | pho-14 | phosphotyrosine protein phosphatase | 20.61 | 0.97 | 22.5 +/-0.4 |
| NCU01889 | 13873 | CC1-PTP | n/a | hypothetical protein | 20.68 | 1.00 | 22.2 +/-0.4 |
| NCU08380 | 20306 | HAD-FCP | csp-6 | plasma membrane phosphatase required for sodium stress response | 20.75 |  | arr |
| NCU03426 | 16425 | CC1-PTP | dsp-1 | pps1 dual specificity phosphatase | 20.76 | 0.95 | 22.5 +/-0.2 |
| NCU03495 | 16430 | PPM | pph-7 | protein phosphatase-7 | 20.77 | 0.98 | 22.2 +/-0.2 |
| NCU03825 | -- | HAD-FCP | n/a | hypothetical protein | 20.78 | 0.94 | nd |
| NCU08158 | 19644 | CC1-PTP | dsp-4 | dual specificity phosphatase | 20.80 | 0.95 | 22.7 +/-0.4 |
| NCU03436 | 11652 and 16003 | PPPL-PPP | tng | serine/threonine-protein phosphatase ppe1 | 20.88 | 0.97 | 22.3 +/-0.3 |
| NCU05049 | -- | CC1-PTP | n/a | hypothetical protein | 20.89 | 0.96 | nd |
| NCU06966 | 14056 | CC3-CDC25 | pty-1 | hypothetical protein | 20.91 | 0.99 | 22.0 +/-0.4 |
| NCU05364 | 12444 | CC1-DSP | pty-3 | tyrosine-protein phosphatase non-receptor type 6 | 20.91 | 0.97 | 22.4 +/-0.3 |
| NCU03246 | 13311 | CC1-PTEN | cdc14 | tyrosine-protein phosphatase CDC14 | 21.03 | 0.98 | 22.3 +/-0.1 |
| NCU01433 | 15790 | PPPL-PPP | ppt-1 | serine/threonine-protein phosphatase 5 | 21.06 | 0.99 | 22.2 +/-0.2 |
| NCU06252 | 14464 | CC1-PTP | dsp-2 | Dual specificity protein phosphatase 2 | 21.08 | 1.01 | 22.0 +/-0.1 |
| NCU01010 | 16679 | CC1-PTP | pty-5 | hypothetical protein | 21.19 | 0.99 | 22.6 +/-0.2 |
| NCU01767 | 12451 | PPM | pph-5 | phosphatase 2C family protein | 21.24 | 0.98 | 22.7 +/-0.5 |
| NCU03333 | 17653 | CC1-PTP | pty-6 | tyrosine-6 | 21.33 | 1.01 | 22.5 +/-0.1 |
| NCU00434 | -- | PPM | n/a | protein phosphatase-8 | 21.38 | 0.97 | nd |
| NCU06330 | 15781 | CC1-PTP | dsp-3 | hypothetical protein | 21.52 | 0.99 | 21.9 +/-0.5 |
| NCU02257 | 16059 | CC1-DSP | pty-2 | protein-tyrosine phosphatase 2 | 21.55 | 0.96 | 22.8 +/-0.1 |
| NCU06563 | 11546 | PPPL-PPP | pp2a (like) | serine/threonine-protein phosphatase PP2A catalytic subunit | 21.77 | 0.98 | fc |
| NCU03114 | 16337 | CC2-SSU72 | rpo-2 | SSU72 | 22.26 | 0.95 | 23.3 +/-0.2 |
| NCU07489 | 11548 | PPPL-PPP | pzl-1 | serine/threonine-protein phosphatase PP-Z | 23.34 | 1.00 | 24.1 +/-0.4 |
| NCU07013 | -- | RTR1 | n/a | hypothetical protein | arr | n/a | nd |
| NCU09300 | n/a - essential | HAD-FCP | fcp-1 | RNA polymerase-24 | -- | -- | -- |
| NCU00043 | n/a - essential | PPPL-PPP | ppp-1 | serine/threonine-protein phosphatase PP1 | -- | -- | -- |
| NCU03804 | n/a - essential | PPPL-PPP | pph-2 | serine/threonine-protein phosphatase 2B catalytic subunit | -- | -- | -- |
| NCU06630 | n/a - essential | PPPL-PPP | pph-1 | serine/threonine-protein phosphatase PP2A catalytic subunit | -- | -- | -- |

-- essential gene
nd not done
arr arrhythmic
fc fails calculation of a valid period in Chrono

**Table 2:** Growth and fertility phenotypes for previously uncharacterized *Neurospora* phosphatases. Phosphatase names, descriptions, phosphatase fold/family, and FGSC reference numbers are similar to Table 1. Growth rate was measured on race tubes and calculated as a percentage of the relevant wildtype strain run at the same time. All strains including wildtype contained the *ras-1*^*bd*^ allele in growth rate assays. Fertility was scored as yes or no, based on the presence of shot ascospores 3-4 weeks after crossing. Asexual growth phenotypes were scored by eye on complete media after 7-10 days of growth at 25°C

| NCU | FGSC origin | Symbol (FungiDb) | Description (FungiDb) | Fold-Family-SubFamily (HMMER) | Growth rate (% of 87-3 ras-1<bd>) | Fertility |  | Asexual growth (description) | Comments |
| --- | --- | --- | --- | --- | --- | --- | --- | --- | --- |
|  |  |  |  |  |  | Female | Male |  |  |
| 01889 | 13873 | -- | hypothetical protein | CC1-PTP | 100 | Yes | Yes | normal |  |
| 05049 | n/a | -- | -- | CC1-PTP | nd | nd | nd | normal | absent from KO collection; ascospores slow to germinate |
| 03825 | n/a | -- | -- | HAD-FCP | nd | nd | nd | normal | absent from KO collection |
| 03731 | 18653 | -- | HAD superfamily hydrolase | HAD-FCP | 90 | Yes | Yes | normal |  |
| 04924 | 20352 | cut-1 | HAD-superfamily hydrolase, variant1 | HAD-NagD-CUT | 95 | Yes | Yes | normal |  |
| 03173 | 18749 | -- | HAD superfamily hydrolase | HAD-NagD-CUT | 99 | Yes | Yes | normal |  |
| 05818 | 20134 | -- | phosphatidyl synthase | HAD-NagD-CUT | 99 | Yes | Yes | normal |  |
| 09683 | 18536 | -- | -- | HAD-NagD-CUT | 95 | Yes | Yes | normal |  |
| 00088 | 16344 | npp-1 | nitrophenylphosphatase-1 | HAD-NagD-PGP | 90 | Yes | Yes | normal |  |
| 01371 | 21200 | -- | UPF0660 protein, variant1 | HAD-NagD-PGP | 57 | Yes | Yes | normal |  |
| 04613 | 15152 | -- | HAD superfamily hydrolase | HAD-NagD-PGP | 95 | Yes | Yes | normal |  |
| 02749 | 19811 | -- | hypothetical protein | PPM | 99 | Yes | Yes | normal |  |
| 00434 | n/a | -- | -- | PPM | nd | nd | nd | reduced macroconidia | absent from KO collection |
| 02150 | 19090 | stp-1 | serine-threonine protein phosphatase-1 | PPPL-PPP | 98 | Yes | Yes | normal |  |
| 07013 | n/a | -- | -- | RTR1 | nd | nd | nd | severely reduced ariel hyphae and macroconidia | absent from KO collection; slow growth |
| 02496 | (16180 x WT) | div-12 | M-phase inducer phosphatase 3 | CC3-CDC25 | nd | yes | nd | nd | Characterized by Ghosh. Reported as infertile for a sexual cross. Has low fertility but does produce homokaryon progeny in my hands. |

nd = not done

### Long period mutants

#### NCU07489 (*pzl-1*), PPPL-PPP family

Deletion of this phosphatase lengthens period of the core clock by ∼2.5hrs at all temperatures, which is the largest period increase identified in our screen (Fig. 3A). Two *pzl-1* orthologs exist in yeast, Ppz1 and Ppz2 (YML016C and YDR436W), due to a duplication event and this phosphatase exists only in fungi; there are no orthologs identified in animals. The PZL-1 catalytic protein dimerizes with one regulatory subunit to form an active phosphatase. Multiple regulatory subunits exist, and these confer substrate specificity for the enzyme. PZL-1 is 64% identical to the much beaer studied catalytic subunit of PP1, PPP-1. PZL-1 regulatory subunits are not annotated in *Neurospora*, and it is not known if the same regulatory subunits can be used by both PZL-1 and PPP-1. Unlike deletion of PZL-1, reduction in PP1 activity is known to shorten period by 1-1.5hrs[15]; therefore, sharing of regulatory subunits seems unlikely.

**Figure 3.**
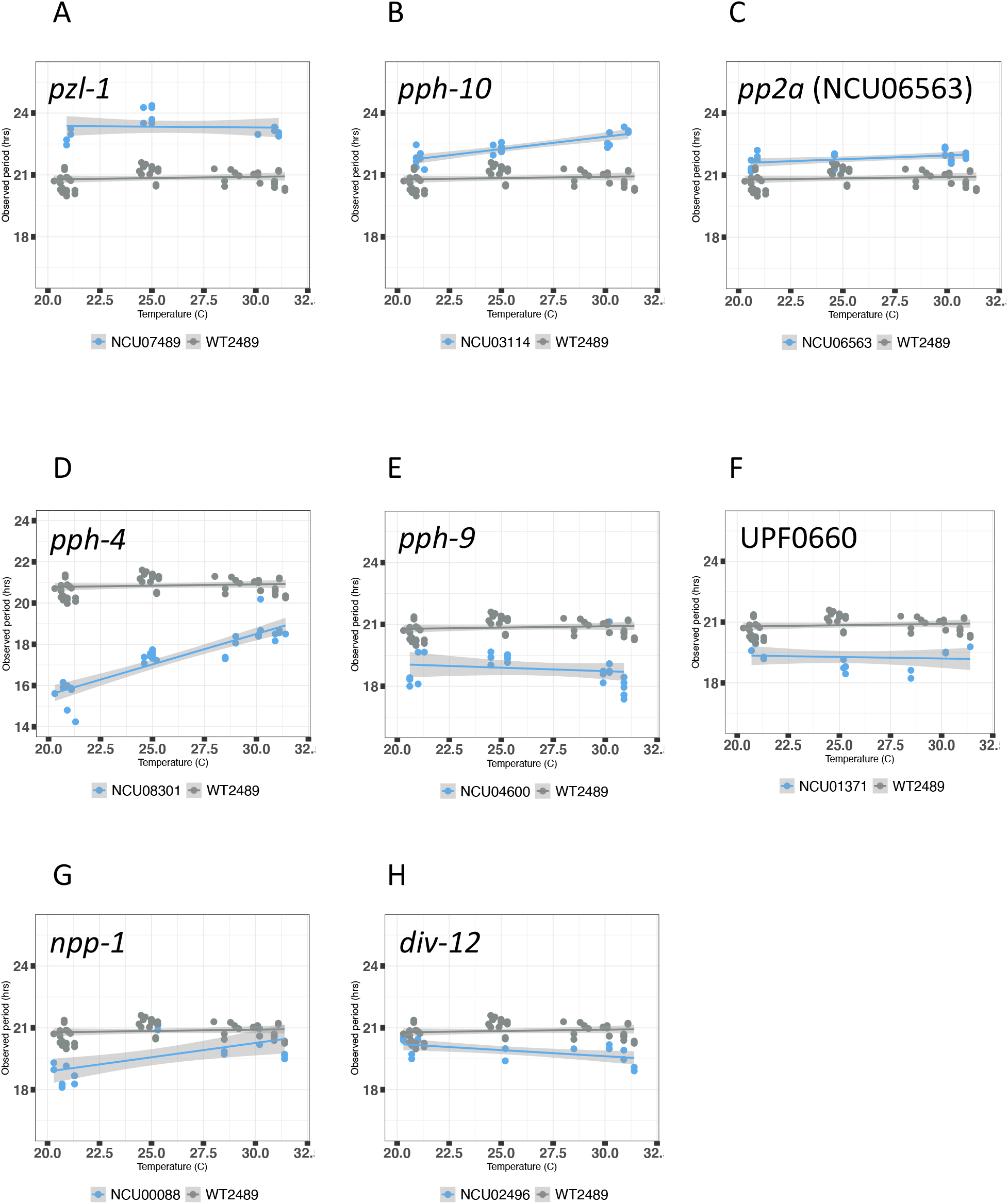
Temperature compensation graphs for selected phosphatase knockouts. (A-H) Gene knockout strains of interest are shown in blue. The composite WT control from all screening runs is shown in gray for comparison. Temperatures ranging from 21°C to 31°C are presented on the x-axis, with corresponding periods graphed on the y-axis. Periods were calculated using cbox-luc reporter signal from at least 2 knockout progeny or transformants tested at 3 temperatures. Best fit regression and 95% confidence interval were calculated using a linear mixed model in R.

#### NCU03114 (*rpo-2*), CC2-SSU72 family

Deletion of *rpo-2* lengthens period by ∼1.4hrs at all temperatures compared to wildtype controls (Fig. 3B). The Ssu72 family is highly conserved from yeast to mammals. It is one of the three related occurrences of a cysteine based catalytic mechanism. Ssu72 has not been studied in *Neurospora*; however, the yeast ortholog (YNL222W) regulates phosphorylation of the RNA pol II C-terminal domain (CTD). The Pol II CTD phosphorylation code controls mRNA transcription initiation, processivity, and termination by altering transcription factors recruited to the polymerase complex. This suggests the longer period of the *rpo-2* knockout could be due to reduced transcription of *frq*, leading to a delay of negative feedback in the clock. This is consistent with prior work that identified BRD-8, a novel auxiliary subunit of the NuA4 histone acetylation complex that also forms a complex with the transcription elongation regulator BYE-1 and whose loss results in a long circadian period[21].

#### NCU06563 (*pp2A*), PPPL-PPP family

NCU06563 is one of two known PP2A catalytic subunits in *Neurospora*. The yeast ortholog is PPG1 (YNR032W) and the phosphatase is conserved to mammals. Care should be taken to avoid confusion with better the studied PP2A catalytic subunit *pph-1* (NCU06630), which is an essential gene sharing 56% identity at the amino acid level. Loss of NCU06563 yields a viable strain but lengthens circadian period by ∼0.9hrs across all temperatures, reduces growth rate, and produces less robust banding on race tubes (Fig. 3C, Fig. 4D). The active PP2A phosphatase is a heterotrimer, consisting of one catalytic, one regulatory, and one scaffolding subunit. Multiple options exist for each subunit of the complex, with substrate specificity achieved through the unique combinatorial assembly. Interestingly, aaempts to reduce function of PPH-1 were either inviable or failed to significantly reduce protein abundance, suggesting PPH-1 and NCU06563 have unique substrates[15]. Two PP2A regulatory subunits (NCU03786 and NCU09377) and one scaffolding subunit (NCU00488) are annotated in *Neurospora*. It is not known which of these interact with NCU06563 for circadian functions, or if there are additional unannotated subunits.

**Figure 4.**
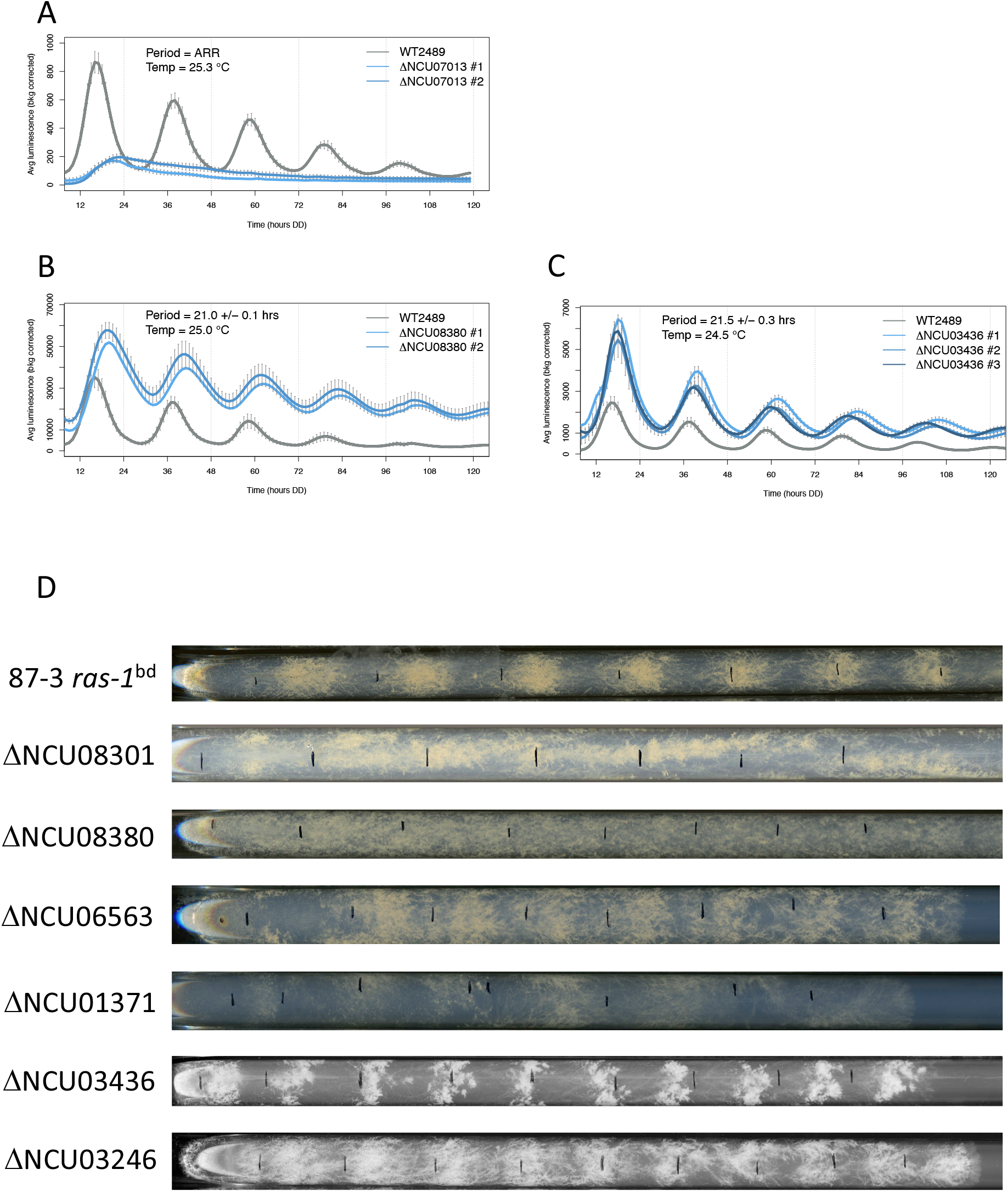
Arrhythmia, phase, and output defects identified by the screen. (A) Knockout of NCU07013 is arrhythmic. Cbox-luc reporter trace at 25°C shown in blue with wildtype comparison in gray. The strain displays an initial light response but does not complete the first circadian cycle. (B) Knockout of NCU08380 (*csp-6*) is phase delayed approximately 4hrs, similar to previous reports. (C) Deletion of NCU03436 (*tng*) is phase delayed approximately 2hrs relative to wildtype control. (D) Output defects as presented by race tubes run at 25°C in constant darkness to observe conidial banding (output) in free-run. One representative race tube shown for each strain. All deletion strains carry the *ras-1*^*bd*^ allele. Arrhythmic output for knockouts of NCU08301 and NCU08380 has been reported previously. NCU06563 (*pp2a*) deletions have weak banding that cannot be accurately scored on race tubes. Deletions of NCU01371 (*upf660)* show an irregular growth rate and lack a regular banding paaern. NCU03436 (*tng*) deletions have normal circadian period, but band appearance is abnormal likely because of developmental defects in the formation of macroconidia. NCU02346 (*cdc14*) deletions show abnormal conidiation in the normal interband region.

### Short period mutants

#### NCU08301 (*pph-4*), PPPL-PPP family

*pph-4* is the only known catalytic subunit of phosphatase PP4 in *Neurospora*. Knockout produced the largest period change of any mutant in our screen, approximately 3.8 hrs short, as well as a temperature compensation defect (Q10 = 0.84) (Fig 3D). Like PP2A, active PP4 is a heterotrimer composed of one catalytic subunit, one regulatory subunit that confers substrate specificity, and one scaffolding substrate which links the catalytic and regulatory proteins. PP4 is highly conserved and present from yeast (YDR075W) to mammals. In most organisms it is an essential gene; however, in *Neurospora* it is dispensable. PP4 regulatory and scaffolding protein(s) are not presently identified in *Neurospora*. A RIP mutant of *pph-4* was previously shown to have a short period circadian phenotype [14, 16]; however, in our present screen period we found the period to be additionally shortened and to have a temperature overcompensation defect as well. We attribute these new observations to nutritional differences caused by a slightly richer solid growth media containing quinic acid (QA). PP4 activity is required for diverse biological processes including the DNA damage response and glucose signaling. In the circadian clock PP4 is hypothesized to act on FRQ to promote protein stability, and on WCC to regulate nuclear/cytoplasmic localization[16].

#### NCU04600 (*pph-9*), PPM fold

*pph-9* deletion shortens period of the core clock by ∼1.9hrs at all temperatures tested (Fig. 3E). Banding output on race tubes is reduced at each successive cycle, although remains rhythmic (Fig. 4D). PPH-9 is a member of the PPM metal-dependent protein phosphatases which rely on a central Mg^+2^ or Mn^+2^ ion to coordinate the catalytic site of the enzyme. Phosphatases in this fold are active as a single subunit, with unique N and C-termini bearing regulatory sequences[22, 23]. PPH-9 is most closely related to PTC2 and PTC3 (YER089C and YBL056W) in yeast, which limit responses to osmostress, downregulate the UPR, and inactivate the DNA damage checkpoint[24-26]. A prior circadian screen in Drosophila identified several possible PPM family phosphatases which lengthened period when lost, but none which shortened it[18].

#### NCU01371 (UPF0660), HAD-NagD-PGP family

Loss of UPF0660 phosphatase shortens the clock period by ∼1.6hrs at all temperatures (Fig. 3F). Rhythmicity is not robust, as evidenced by reporter traces which deviate from a smooth sine wave (data not shown). Race tubes demonstrate impaired and irregular growth rate with lack of normal circadian output banding (Fig. 4D). The UPF0660 phosphatase is homologous to yeast GEP4 (YHR100C), also known as PGP phosphatase. This enzyme is present in fungi and plants; however, orthologs are absent in mammals. The UPF0660 phospholipase is found in the mitochondrial matrix space where it dephosphorylates phosphatidylglycerolphosphate to produce phosphatidylglycerol[27]. Loss of this phosphatase is expected to alter mitochondrial functions, although it is unclear why this might shorten the period of the clock.

#### NCU00088 (npp-1), HAD-NagD-PGP family

Deletion of *npp-1* phosphatase shortens period by ∼1.3hrs at 25C (Fig. 3G) but maintains normal clock output. The knockout shows a slight overcompensation with a Q10 value of 0.93 (Table1). The yeast ortholog PHO13 (YDL236W) is found in the nucleus and the cytoplasm, where it functions as a metabolic repair enzyme. In yeast the loss of PHO13 alters metabolism by upregulating the pentose phosphate pathway. This is most studied for the ability to increase xylose metabolism for biofuels with yeast.

#### NCU02496 (*div-12*), CC3-CDC25 family

Knockout of *div-12* phosphatase shortened period by ∼0.9hrs. The yeast ortholog MIH1 (YMR036C) is a protein tyrosine phosphatase that is best known for opposing the activity of Swe1 kinase to control the activity of cell cycle regulator Cdk1[28]. MIH1 is regulated by activity of casein kinase 1 (CK1) and dephosphorylated by pp2A[29], both of which are known to participate in maintenance of proper circadian timing. In yeast MIH1 also regulates the mitotic spindle and acts in endocytosis and plasma membrane recycling.

### Arrhythmic mutants

#### NCU07013 (hypotheUcal protein), RTR fold

NCU07013 is the only member of the RTR fold present in *Neurospora*. Deletion strains are viable but lack a core clock circadian rhythm (Fig. 4A), show reduced growth rate, and are defective in aerial hyphae and production of macroconidia. At the level of the core clock, we observe an initial rise in reporter signal after the light-to-dark transition, indicating the strain remains viable with intact light sensing at the start of our assay. At the highest temperature tested an occasional technical replicate will oscillate briefly for 1-2 cycles; however, this is rare and not reproducible (data not shown). NCU07013 has not been studied in *Neurospora*, but the yeast homologs, RTR1 and RTR2 (YER139C and YDR066C), regulate RNA pol II through dephosphorylation of the C-terminal domain (CTD). Pol II associated factors are recruited based on a CTD phosphorylation code that dictates polymerase initiation, processivity, and termination[30]. The human ortholog RPAP2 also appears to regulate Pol II activity. We predict arrhythmicity of the NCU07013 knockout is caused by global downregulation of RNA synthesis and mis-regulation of protein abundance across many biological pathways.

### Altered phase or output

We identify two phosphatase knockouts, *csp-6* (NCU08380) and *tng* (NCU03436) that exhibit normal circadian period, but are phase delayed. The *csp-6* phase delay was previously reported by our lab[31]. Rhythmic circadian output was absent or severely affected in knockouts of *pph-4* and *csp-6* (Fig4D), matching previously published data[16, 31]. We observed reduced banding in knockouts of *pp2a* and UPF660 (Fig. 4D). Banding output is present for knockouts of *tng* and *cdc14* but is abnormal in appearance (Fig. 4D). This is likely due to developmental defects of these strains which affect growth rate and production of conidia.

#### Non-circadian phenotypes

We performed basic phenotypic assessment of deletion mutants of 11 non-essential phosphatases from the *Neurospora* knockout collection that were not previously characterized, plus four novel knockout strains constructed for this screen.

Asexual growth on complete media slants was altered in knockouts of NCU00434 and NCU07013. The NCU00434 deletion strain produced aerial hyphae but showed greatly reduced macroconidiation. Knockouts of NCU07013 lacked both aerial hyphae and macroconidia and grew more slowly than wildtype controls. Interestingly the knockout of NCU01371 exhibited normal growth and conidiation on complete media, in contrast to the reduced/variable linear growth rate observed on race tubes containing minimal media.

Two deletion mutants displayed alterations in the sexual phase of the *Neurospora* lifecycle. Ascospores from a cross between wildtype and a NCU05049 heterokaryon yielded viable homokaryotic progeny, although these germinated more slowly than wildtype siblings. For unknown reasons homokaryons could not be isolated from all crosses. NCU02496 (FGSC16180) was previously characterized and reported to be sterile in a sexual cross. We were able to generate homokaryotic progeny when this knockout was used as the female parent, albeit at a very low frequency.

Linear growth rate was similar to the wildtype control for all knockouts tested, with the exception of NCU01371 as already discussed.

## DISCUSSION

We were surprised to find few phosphatase knockouts that altered period and only one resulting in arrhythmicity. Combining our results with published data suggests that many phosphatases acting on core circadian clock components are clustered in the PPPL-PPP family. These phosphatases are believed to have ancient origins dating back about four billion years to the last universal common ancestor and are highly conserved across species. This is consistent with established phosphorylation machinery being repurposed to regulate circadian clocks, the earliest of which evolved approximately 2.5 billion years ago.

Interrogating specific PPPL phosphatases through chemical inhibition is challenging due to the high degree of conservation in the catalytic site between members of the fold. In this screen we employed knockouts of PPPL catalytic subunits to target individual phosphatases without confounding effects. A logical next step for examining clock-relevant substrates of these phosphatases is to identify the regulatory/scaffolding subunits conferring substrate specificity, and further screen for circadian functions. Clock components are extensively phosphorylated in a time-dependent pattern that is not well understood. It is possible that activity of one phosphatase catalytic subunit may both shorten and lengthen period, depending on which regulatory subunits are used. Adding additional complication is the fact that some regulatory/scaffolding subunits are shared between closely related catalytic subunits and therefore could alter the activity of multiple phosphatases simultaneously.

Functionally, several of the phosphatases altering period length are known to regulate RNA Pol II (RNAPII) C-terminal tail (CTD) phosphorylation. The CTD phosphorylation code recruits transcription, elongation and termination factors to the polymerase complex during production of an mRNA. Because the *Neurospora* clock is a TTFL, altering transcriptional efficiency of the negative regulator *frq* is likely to impact function or result in an arrhythmic clock. RNAPII CTD regulation has broad effects on mRNA abundance, therefore interpretation may be confounded by wide-ranging changes in protein levels[32].

### LIMITATIONS

We relied on the *Neurospora* knockout collection as the starting point for our screen.While these strains were thoroughly validated during construction, a collection of this size almost certainly contains errors or unknown additional mutations. As with any screen, our results are limited by the selection of gene knockouts phenotyped. We intentionally did not phenotype families or subfamilies thought to exclusively target non-protein substrates. Given the many effects of PPPL fold phosphatases identified, perhaps the PPPL-PAP family should be screened in the future.

We attribute circadian phenotypes observed in the knockouts to loss of phosphatase function. These mutants were designed as loss of the entire open reading frame (ORF) of the targeted protein, not simple inactivation of enzymatic function. It is possible, although less likely, that loss of a non-catalytic domain within the protein is responsible for the observed circadian behavior.

## MATERIALS AND METHODS

Knockout strains were obtained from the Neurospora knockout collection or constructed using similar methodology[33, 34]. The cbox-luc reporter was integrated into each knockout by transformation or crossing[35]. The *ras-1*^*bd*^ mutation was integrated by crossing with the stock strain 87-3 *ras-1*^*bd*^.

Monitoring of luciferase reporter activity was performed as described previously, with minor modifications to the media. Our solid media contained 1X Vogel’s salts, 0.03% glucose, 0.01M quinic acid ,0.05% arginine, biotin, luciferin and 1.5% agar. Rhythms were monitored for at least 5 days under constant conditions. For temperature compensation we selected target temperatures of 21°C, 25°C and 29°C. The actual temperature was monitored during each run and plotted during data analysis.

Race tubes were run on solid agar media at 25°C in constant darkness as previously described. Growth fronts were marked every 24 hours, and analysis of period was performed using the ChronOS software. Growth rate was computed as a percentage of the wildtype run on the same day.

Sexual crosses were performed against wildtype strains 2489 or 328-4 using Westagaard crossing media. A cross was scored as fertile if ascospores were shot onto the tube or lid 3-4 weeks after mating. Ascospores produced were not assessed for viability.

Growth and conidiation on complete media were assessed on slants grown at 25°C under constant light for 7 days. Slants were observed visually for height of ariel hyphae, production of conidia, and color.

## FIGURE LEGENDS

**Supplementary Figure 1.**
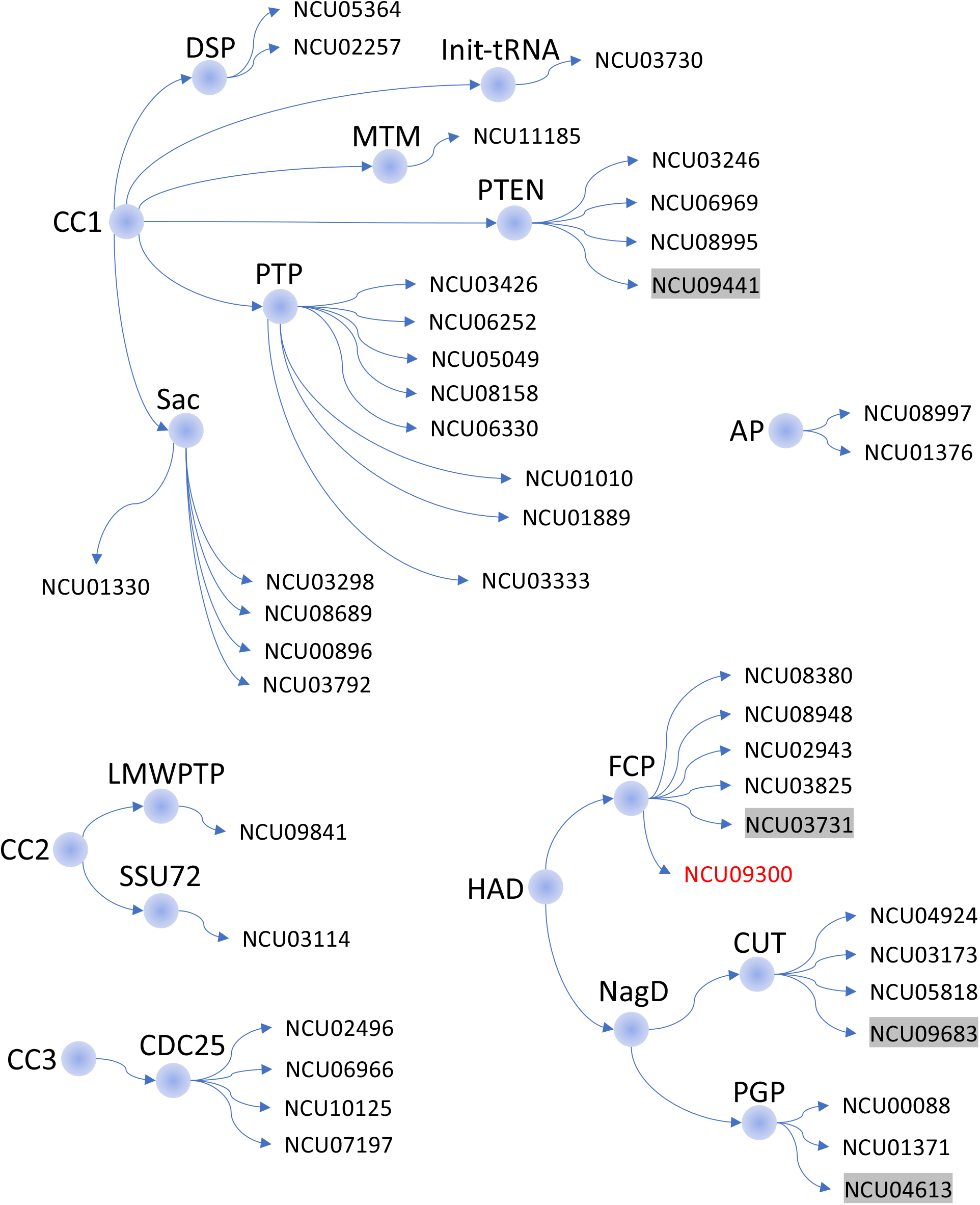
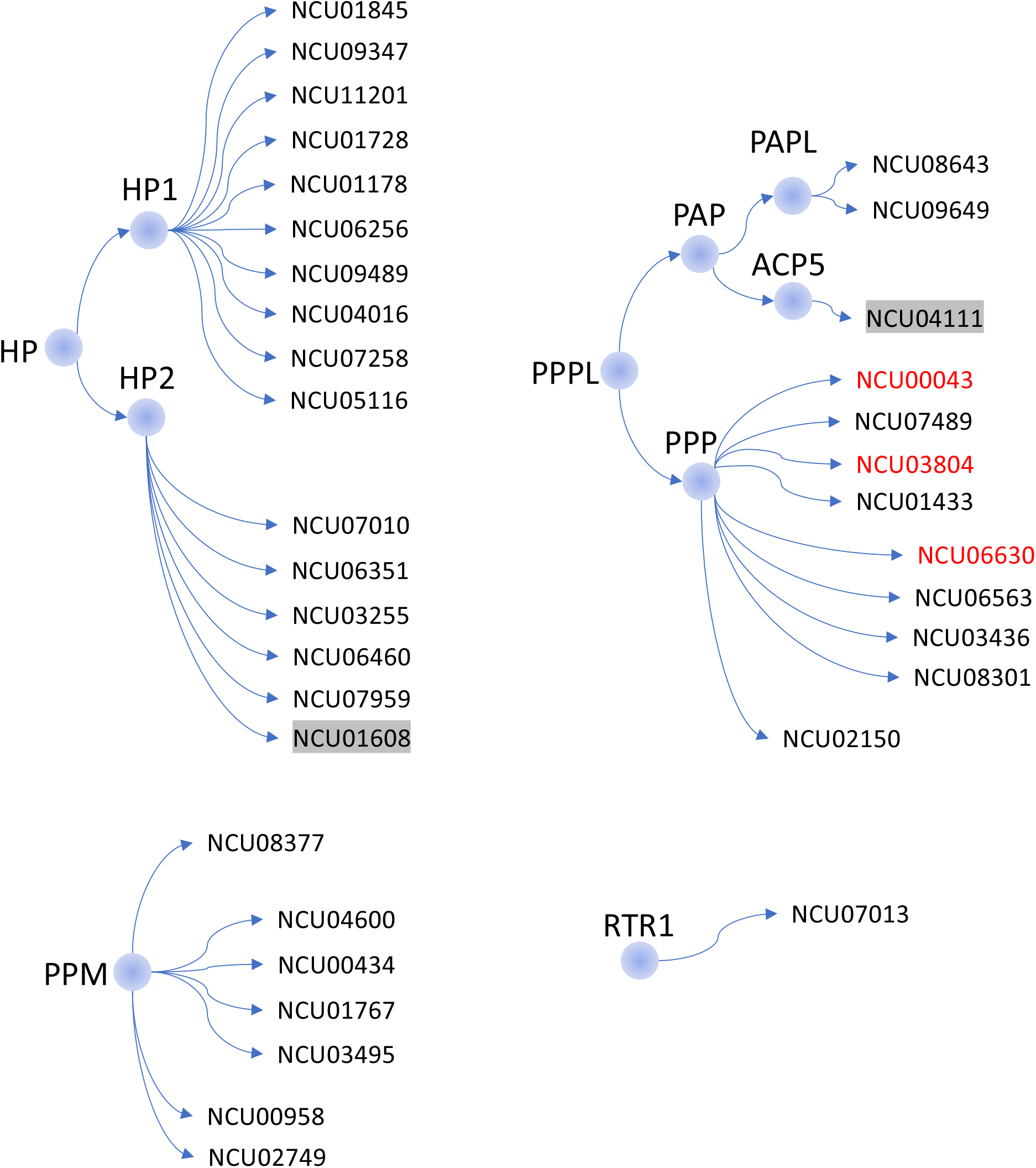
Related to Figure 2. All *Neurospora* phosphatases identified by HMMER. Phosphatases are arranged by fold and family, similar to Figure 2. Non-protein phosphatases and pseudophosphatases are included.

**Supplementary Table 1:** Original HMMER phosphatase profile search results for the *Neurospora crassa* genome.

## ACKNOWLEDGEMENTS

We thank Dunlap-Loros lab members for helpful discussions. This work was supported by NIH R35GM118021 to JCD. The authors declare no conflicts of interest.

